# Estimation of cell cycle kinetics in higher plant root meristem links organ position with cellular fate and chromatin structure

**DOI:** 10.1101/2021.01.01.425043

**Authors:** Taras Pasternak, Stefan Kircher, Klaus Palme

## Abstract

Plant root development is a complex spatial-temporal process that originates in the root apical meristem (RAM). To shape the organ’s structure signaling between the different cells and cell files must be highly coordinated. Thereby, diverging kinetics of chromatin remodeling and cell growth in these files need to be integrated and balanced by differential cell growth and local differences in cell proliferation frequency. Understanding the local differences in cell cycle duration in the RAM and its correlation with chromatin organization is crucial to build a holistic view on the different regulatory processes and requires a quantitative estimation of the chromatin geometry and underlying mitotic cell cycle phases’ timing at every cell file and every position. Unfortunately, so far precise methods for such analysis are missing.

This study presents a robust and straightforward pipeline to determine in parallel the duration of cell cycle’s key stages in all cell layers of a plant’s root and their nuclei organization. The methods combine marker-free techniques based on the detection of the nucleus, deep analysis of the chromatin phase transition, incorporation of 5-ethynyl-2′-deoxyuridine (EdU), and mitosis with a deep-resolution plant phenotyping platform to analyze all key cell cycle events’ kinetics.

In the *Arabidopsis thaliana* L. RAM S-phase duration was found to be as short as 20-30 minutes in all cell files. The subsequent G2-phase duration however depends on the cell type/position and varies from 3.5 hours in the pericycle to more than 4.5 hours in the epidermis. Overall, S+G2+M duration in Arabidopsis under our condition is 4 hours in the pericycle and up to 5.5 hours in the epidermis.

Endocycle duration was determined as the time required to achieve 100% EdU index in the transition zone and estimated to be in the range of 3-4 hours.

Besides Arabidopsis, we show that the presented technique is applicable also to root tips of other dicot and monocot plants (tobacco (*Nicotiana tabacum* L.), tomato (*Lycopersicon esculentum* L.) and wheat (*Triticum aestivum* L.).

## Introduction

The duration of the mitotic cell cycle as well as endo-cycle, including its phases, are essential characteristics for harmonized organ growth kinetics. Investigation of cell cycle duration in plants was first done in 1951 (Brown, 1951) in pea (*Pisum sativum* L.) roots by quantification of the ratio of cells in a certain stage of the cell cycle to the total number of meristematic cells. The interphase duration was found to be 23 hours, prophase 2 hours, metaphase 25 minutes, anaphase 5 minutes, and telophase 22 minutes. Thereafter cell cycle duration was investigated by various different methods. Clowes (1961) as well as Van’t Hof and Sparrow (1963) proposed a method based on H^3^-thymidine incorporation into the replicating DNA. Van’t Hof (1967) further modified this method by additional colchicine treatment in order to accumulate cells in mitosis for better quantification. Using this approach, the entire cell cycle duration was found to be 24 hours in pea (*Pisum sativum* L.), sunflowers (*Helianthus annuus* L.), and bean (*Vicia faba* L.). Recent studies used 5-Ethynyl-2’-deoxyuridine (EdU) as non-radioactive, fluorescent alternative to H^3^-thymidine incorpration (Buck et al., 2008) enabling for example Hayashi et al. (2013) to determine a time window of 17 h for Arabidopsis cell cycle duration during root meristem proliferation. Some researchers estimated the duration of the different cell cycle phases G1, S, G2, M. Using the Van’t Hof method, the S-phase duration in pea (*Pisum sativum* L.), sunflowers (*Helianthus annuus* L.) and bean (*Vicia faba* L.) was found to be 4.5 hours (i.e. 30% of total cell cycle length) (Van’t Hof, 1969). More recently, a combination of EdU pulse labelling with flow cytometry (Mickelson-Young et al., 2016) estimated S-phase duration in Arabidopsis to be 2-3 hours.

Besides the use of labeled nucleotide analogs, several other “kinematic” methods were developed to investigate cell cycle duration in the Arabidopsis root (Beemster and Baskin, 1998; Fiorani and Beemster, 2006). Newly developed stripflow software combined with kinematic data acquisition gave a cell cycle duration in one of the root tissues (cortex) of 14,7±0,9 hours (Yang et al., 2017). A similar Rate of Cell Production (RCP) method (Ivanov and Dubrovsky, 1997) determined a cell cycle duration of 11-13 h for cortex cells of different lines of *Pisum sativum*. Another recent approach to investigate cell cycle duration is the use of marker lines with a dual-color marker system (Yin K., et al., 2014). With this method cell cycle duration was found to be about 16 hours in root epidermis cells, including 3 hours of S-phase. To simplify such an approach a combination of markers was introduced in a single construct to simultaneously label all cell cycle stages and allow to detect cell cycle kinetics in living roots (Desvoyes et al., 2020). However, similarly as in Yin K. et al. (2014), these lines are generated for Arabidopsis only, and require costly efforts for mutant line analysis or transfer to other species.

The kinematic method’s main drawback is that it applies mainly to the outer cell layers (epidermis/cortex) and is based on the assumption that cell cycle duration is constant throughout the proliferation zone. But it′s well-known since 1961 (Clowes, 1961) that cell cycle duration depends on cell position and cell type. For example, it was reported that in *Convolvulus arvensis* cell cycle duration in the central cylinder and cortex were significantly different (21 h and 27 h, respectively), while in the quiescent center, it reaches even 420 h (Phillips & Torrey,, 1972). However, such differences have not been addressed in most of the published methods. Additionally, it was only recently shown that cell cycle duration can be regulated by H3 histone modification and was different in the proximal and distal zone of Arabidopsis root epidermis (Otero et al., 2016).

In summary, many fundamental questions about positional differences in the duration of the cell cycle phase in functionally distinct tissues and cell files and its potential link with nucleus landscape in the root meristem remain unclear. Our working hypothesis is that different cell files may exhibit different cell cycle duration and kinetics dependent on chromatin geometry. These differences may aim to compensate for the differences in cell growth kinetics of the different cell lines in the context of a compact organ. To resolve this issue, the precise estimation of chromatin geometry and cell cycle kinetics in each cell in the frame of an organ coordinate system is required. Here we provide a simple and robust method for determination of S-phase, G2-phase, and M-phase duration in roots of higher plants and demonstrate its value. The proposed method does not require marker lines, allows to determine the duration of S, G2, and M stages in all cell files simultaneously and independently and is therefore generally applicable in plants. A combination of EdU incorporation, mitosis labeling and nuclear geometry analysis with an accurate coordinate system (iRoCS) (Schmidt et al., 2014; Pasternak et al., 2021) allowed us to generate precise maps of main cell cycle events in the root of Arabidopsis as well as several other plants.

## Results

### Divergence of cellular parameters in different tissues and cell files of the Arabidopsis RAM

The Arabidopsis root apical meristem (RAM) has a radial structure built up by functionally distinct cell layers. In order to perform a high/resolution analysis of the Arabidopsis root we first used the iRoCS analysis pipeline for a detailed analysis of the geometry of all cells within this organ and nucleus morphology in these cells (Schmidt et al., 2014; Pasternak et al., 2021).

We demonstrated a strong divergence between and within different cell types regarding chromatin landscape (Figure 1) and cell volume (Figure S1). Interestingly, epidermis and cortex cells exhibit a significant cell volume increase even in the proliferation domain, while for the pericycle and endodermis cells, this tendency is not visible (Figure S1). The chromatin structure also differs significantly among cells even in the proliferation domain with a strong divergence of the volume of the nuclei at the end of the proliferation zone between outer (cortex/epidermis) and inner cell layers (Figure 1). Next, we compared individual nuclei of atrichoblast precursor (AT) and pericycle cells and demonstrated that the total volume of AT nuclei is around 200 µm^3^ while the volume occupied by DNA is about 120 µm^3^. At the same time, these parameters for pericycle cells are approximately 50 and 40 µm^3^ correspondingly. However, in both cases, total volume of condensed chromatin during mitosis is around 20 µm^3^. In consequence, the magnitude of chromatin packing is about a factor of six for atrichoblasts but only about two for pericycle cells. The interchangeable morphology of nuclei usually has not been considered as a potential player in cell cycle transition (Skinner et al., 2017). Our results strongly suggest that nuclear shape has an effect on cell cycle transition because the process is accompanied by chromatin phase changes. However, different cell types exhibit different nuclear shapes/size (Fig. 1). Cell cycle transitions can be considered as changes in the nuclear organization: nucleosomes and chromatin disassemble during replication, which is required to relax chromatin structure, and oppositely, chromatin condenses to the chromosomes during mitosis (Pecinka et al., 2020). Significant differences in nucleus structure between cell files led us to hypothesize that cell cycle duration (G1 and G2) may differ in different cell files and even vary therein at different positions in the root to compensate for variations in cell size and could be also dependent on chromatin remodeling during mitosis and DNA replication.

**Figure 1.**
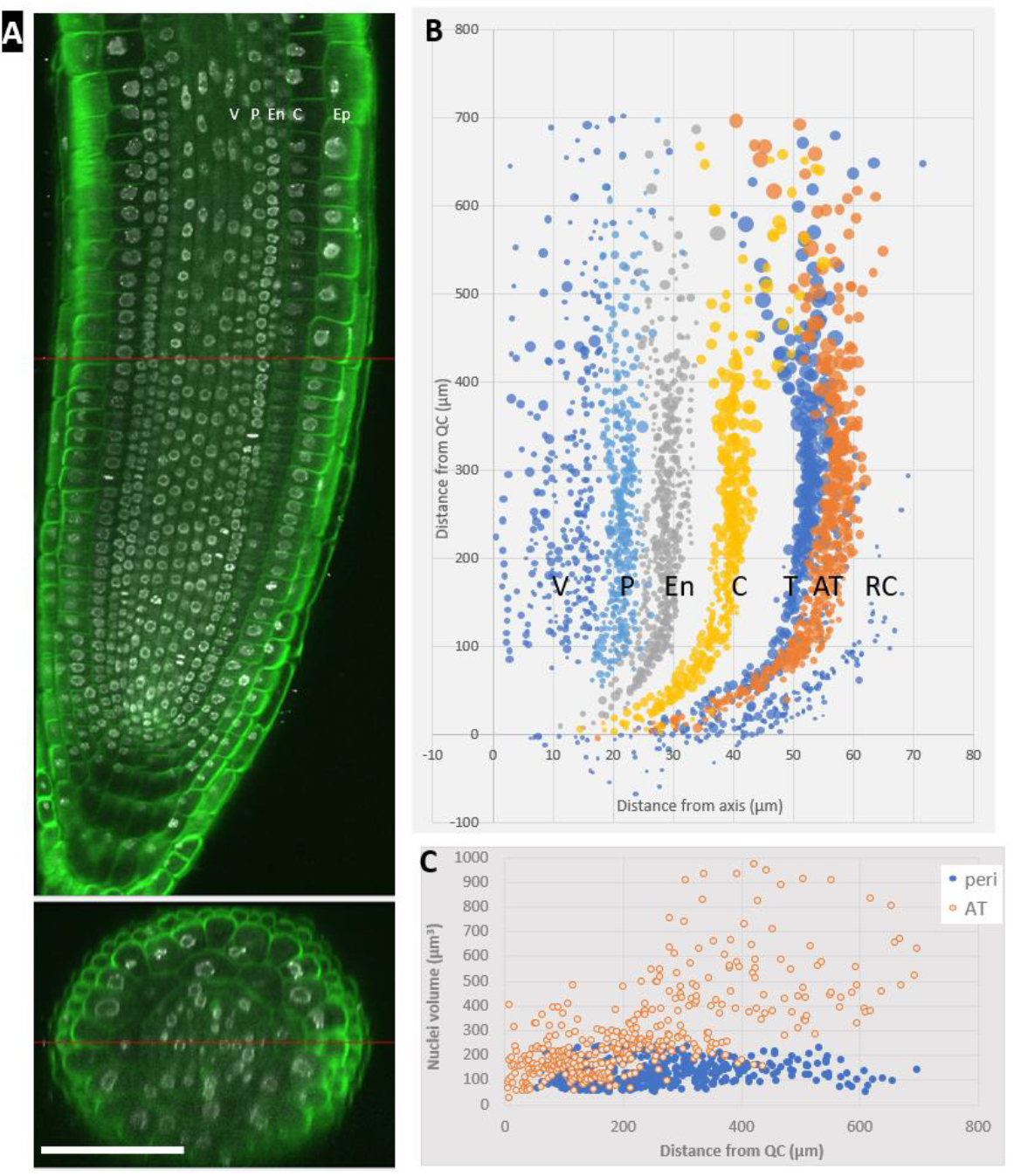
Divergence of nucleus geometry in the proliferation domain of Arabidopsis root. (A) Nuclear landscape after DAPI labelling. Roots were analysed five days after germination (DAG). V-vasculature; P – pericycle; En – endodermis; C - cortex; Ep-epidermis; (B) – Map of nuclear volume; Each bubble represents individual nuclei, radius of the bubbles reflect relative nucleus volume; V-vasculature; P – pericycle; En – endodermis; C – cortex; T-trichoblast precursor; AT – atrichoblast precursor; RC-root caps. (C) – evaluation of the nucleus volume in atrichoblast (orange) and pericycle (blue). Scale bar: 50 µm.

**Figure S1.**
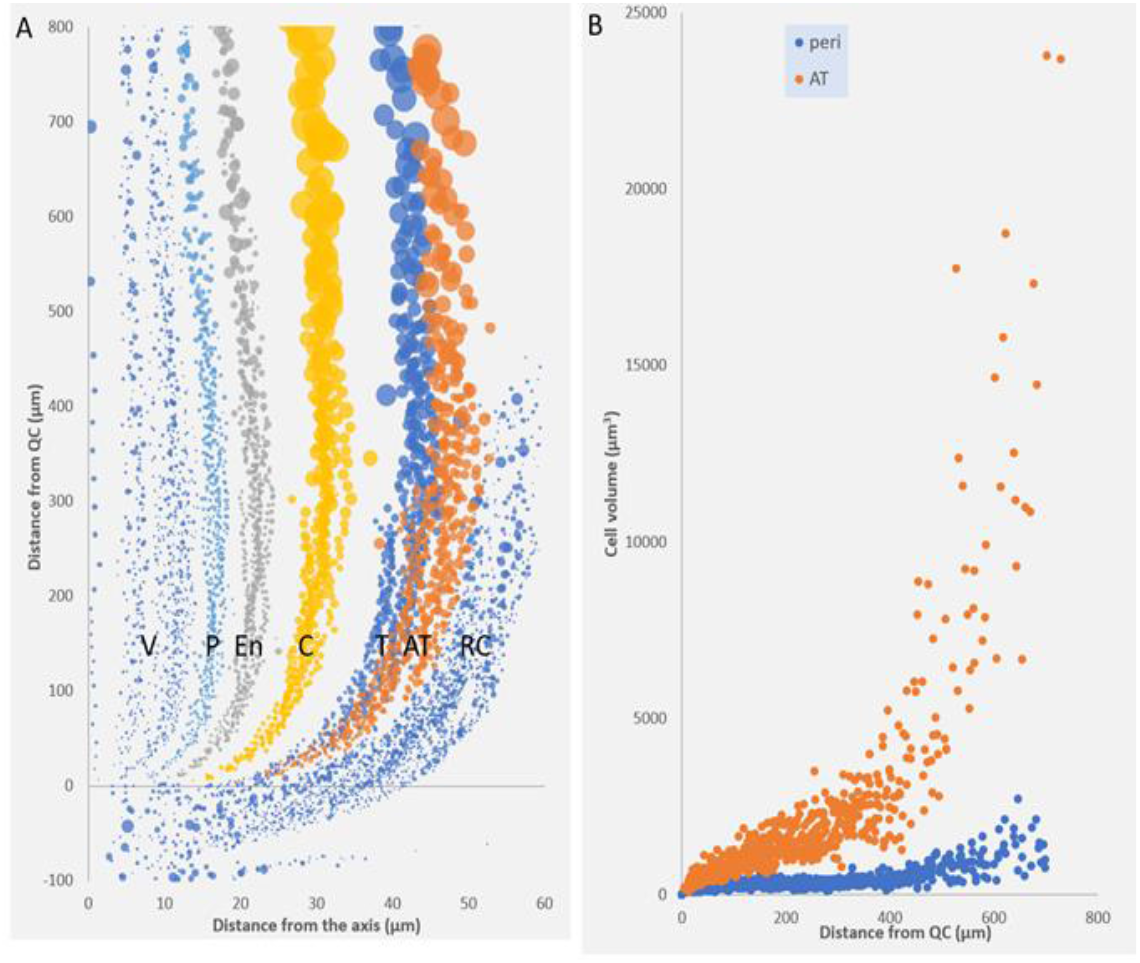
Volumetric cell analysis. (A)-overview of the cell volume as function of the radial distance and distance from QC; Roots were analysed five days after germination (DAG). Each bubble represents an individual cell, radius of the bubble reflect relative cell volume. V - vasculature; P-pericycle; En-endodermis; C-cortex; T-trichoblast precursor; AT-Atrichoblast precursor; RC-root cap. (B) AT and pericycle cell analysis.

### Work-flow to determine cell cycle parameters

To determine the duration of the different cell cycle phases we used the workflow presented in Figure 3. To minimize potential variations, we performed the experiments by simultaneous determination of all stages in a single seedling population: after 4,5 days growth under standard conditions Arabidopsis seedlings were distributed to 6 vials, EdU have been added at the given time points to monitor DNA replication and all samples were fixed in the end of the time course simultaneously.

**Figure 2.**
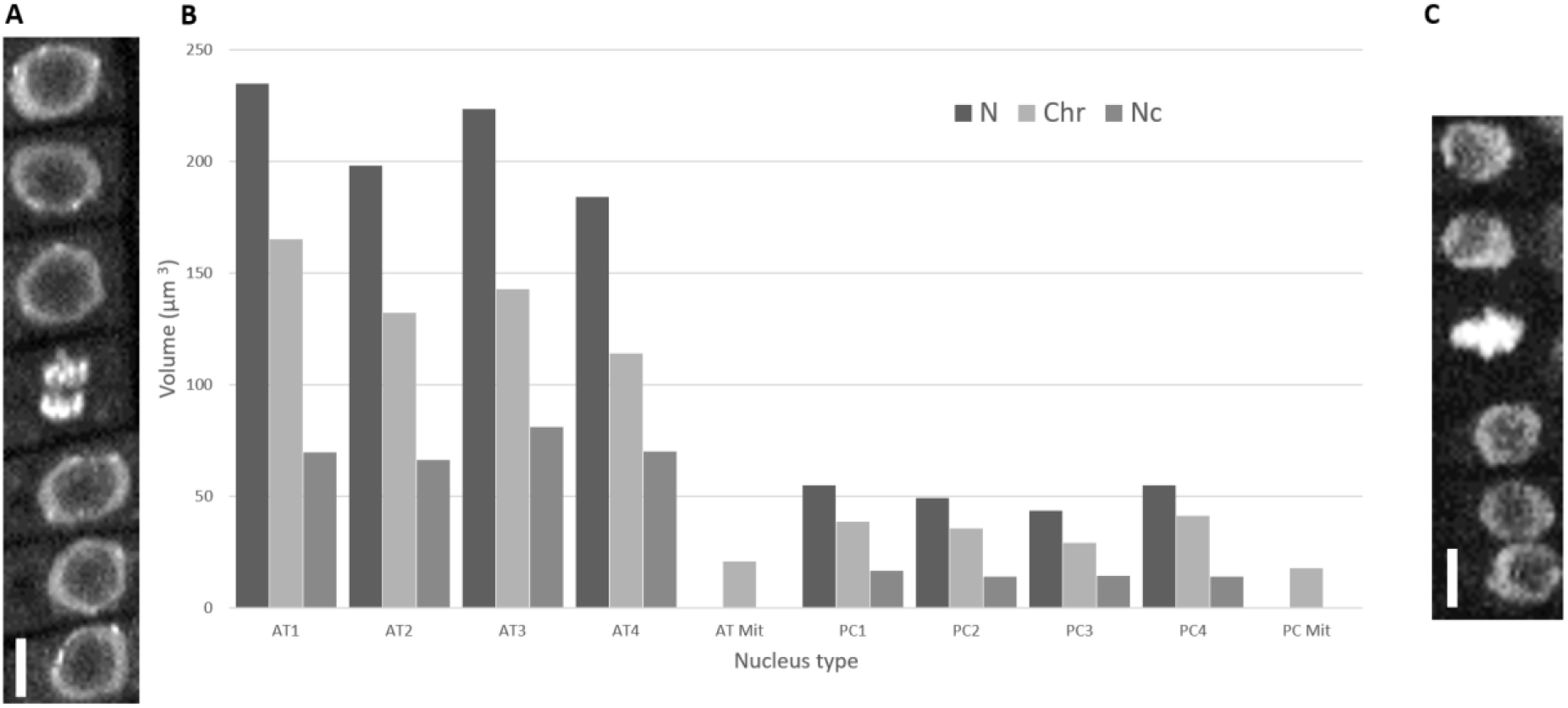
Feature of the individual nuclei of atrichoblast and pericycle cells. (A) Image of Atrichoblast nuclei; (B) Quantitative analysis of the nuclei parameters (N-nuclei volume; Chr-chromatin volume; Nc-nucleoli volume), AT-atrichoblast nuclei 1-4; PC-pericycle nuclei 1-4; (C) image of the pericycle nucleus. AT1 - AT4 and PC1 - PC4 mean individual nuclei. Scale bar – 5 µm.

**Figure 3.**
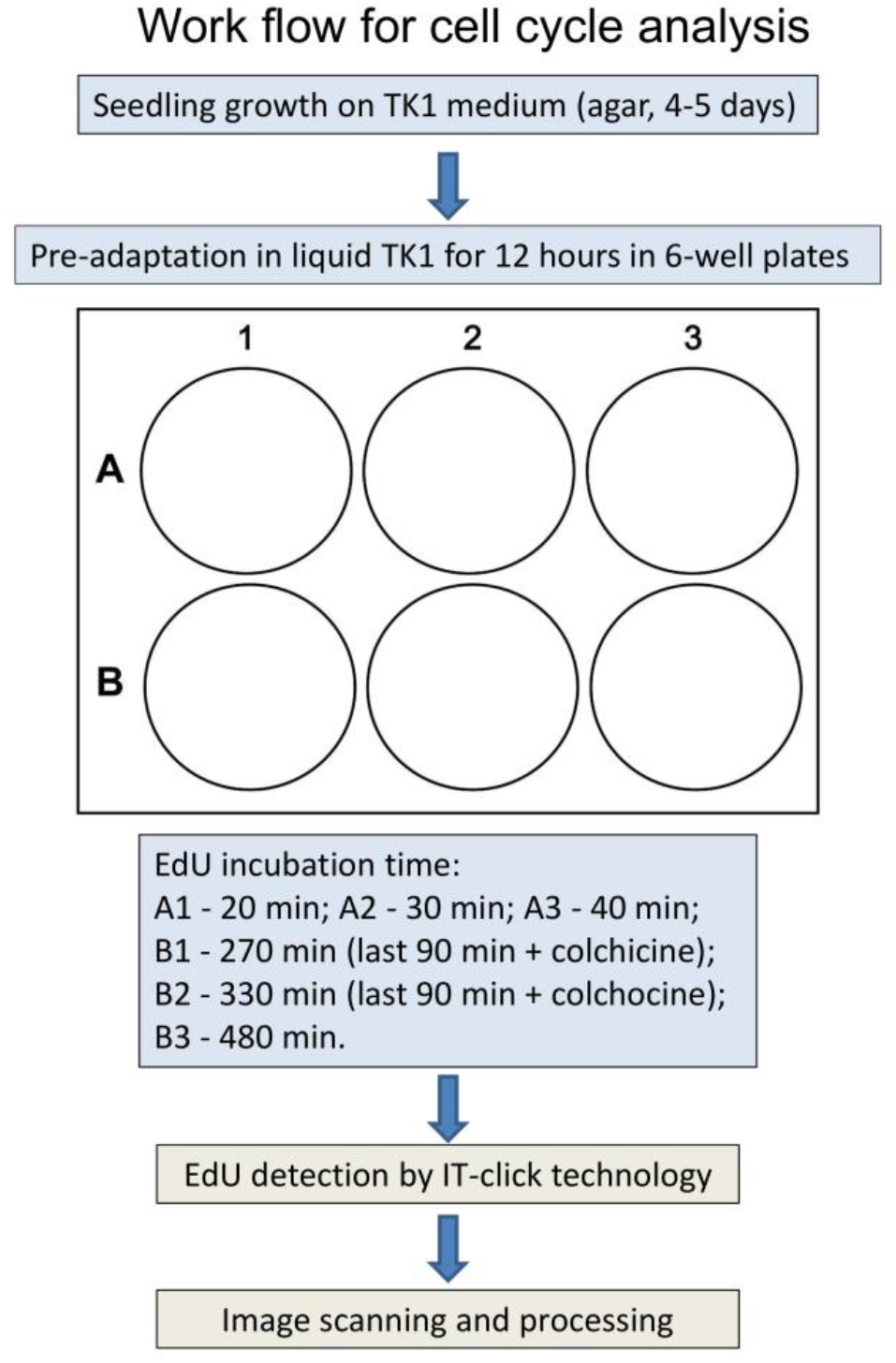
Workflow for estimation of cell cycle phases duration in Arabidopsis.

### S-phase duration

Using this work-flow (Fig. 3) we determined the S-phase duration by estimating the minimal time required for full co-localization of DNA and EdU derived signals. After incubation for 20 min, 30-40% of the total number of the nuclei showed complete EdU and DAPI co-localization (Figure 4). Interestingly, after 20 min of incubation all cell files, even most central stele tissues in the Arabidopsis root tip showed many cells performing DNA replication. From these data we conclude that the S-phase lasts about 20-30 min. We note that in dense chromatin (heterochromatin) EdU (Figure 3, blue arrows) incorporated faster compared to eu-chromatin, but the later cells are mainly located in the transition domain where cell elongation is detectable (post-mitotic domain).

**Figure 4.**
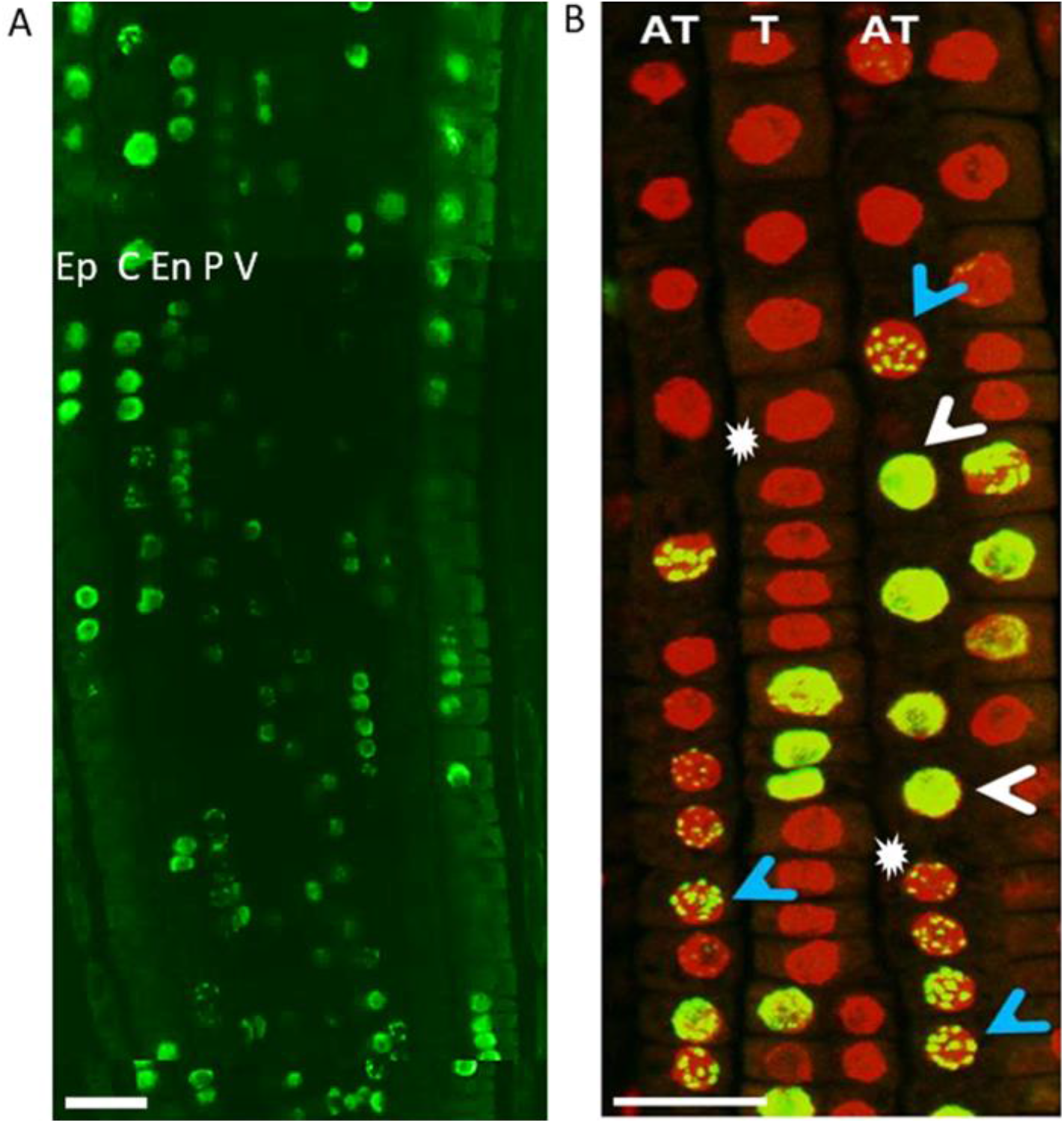
DNA replication in Arabidopsis root after 20 min. of EdU incubation. (A) Maximum projection of the epidermal cell layer of a representative root is shown. DAPI staining is presented in red, EdU signals in green. (B) overview of the distal RAM zone. White arrow heads point to examples of entire labelling, blue arrow heads mark partial labelling of only condensed DNA spots. The white asterisks mark visible cell elongation in the trichoblast (T) and atrichoblast (AT) cell files (onset of transition zone). V - vasculature; P - pericycle; En-endodermis; C - Cortex; Ep - epidermis. EdU-positive nuclei are in green. Scale bar: 20 µm.

### G2 duration

To determine the duration of the G2 phase, we first analyzed the minimal time needed for appearance of EdU-positive cells’ in the mitotic stage. For this purpose, EdU incubation was performed for 90, 180, 210, 240 and 270 minutes. EdU-positive cells passing through mitosis (i.e. pass DNA replication and G2 phase) in the pericycle and endodermis were detected after 240 minutes, but not after 210 minutes. The first EdU-positive mitotic cells in the cortex and epidermis were detected after 270 minutes. Based on this observation, we chose EdU incubation up to 270 minutes for detailed analysis and then quantified EdU-positive mitotic cells’ ratio in each cell file. To prevent EdU-incorporating cells from passing through mitosis and cell division, colchicine was added after 180 minutes of EdU incubation.

This procedure allowed us to build a detailed map of the mitosis distribution from a typical root (Figure 5A). This map was built on calculated average values of about 6000 individual cells obtained from three independent roots (Figure 5B). From these data we conclude that in the inner layers (endodermis, pericycle, and vasculature) cells passing the (S+G2) phase in 180 to 270 minutes and are distributed uniformly in the whole proliferation domain. However, in the cortex and epidermis, only few cells located close to the QC passed the (S+G2) phase during this time span, while more distally located cells obviously exhibit a longer G2 duration.

**Figure 5.**
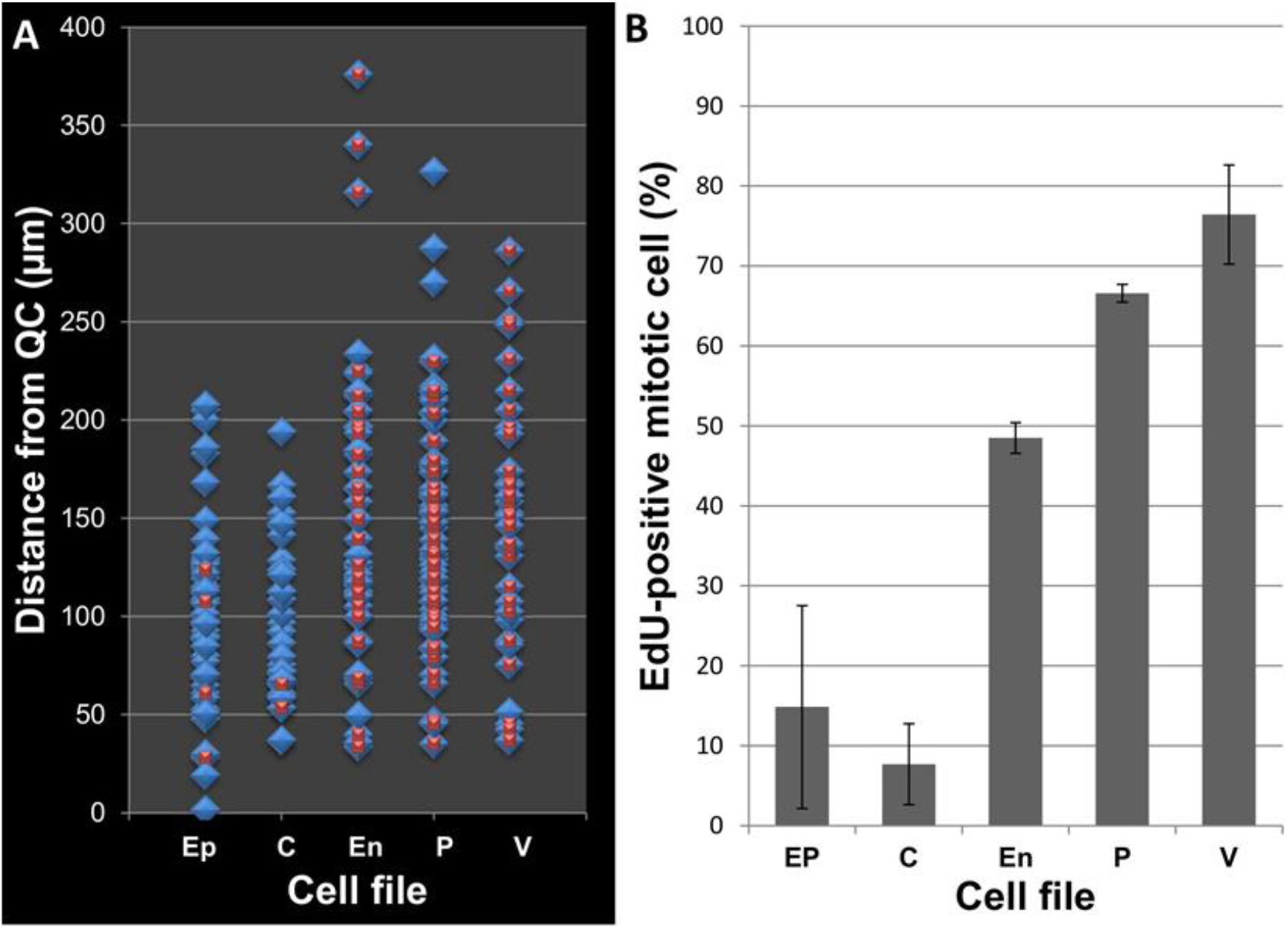
Vertical and horizontal gradients of G2 duration. (A) Mitosis distribution in different tissue layers of 4.5 days old Arabidopsis root tips. Seedlings were incubated in the presence of EdU for 180 minutes, subsequently colchicine was added for further 90 minutes. Seedlings were fixed immediately afterwards and subjected to analysis. Mitosis events only are depicted in blue; mitotic cells exhibiting EdU incorporation are marked blue plus red. (B) Average percent of EdU-positive mitotic cells in different cell files. Ep - epidermis, C - cortex, En - endodermis, P - pericycle, V - vasculature.

To further clarify G2 duration, we increased EdU incubation time to 330 minutes and added colchicine for the last 90 minutes. Results of a typical root analysis demonstrated that the number of EdU-positive cells entering in mitosis after that time period increased to 80% in the cortex/epidermis, while in more inner cell layers, we observe 90-100% of EdU-positive mitosis (Figure S2).

**Figure S2.**
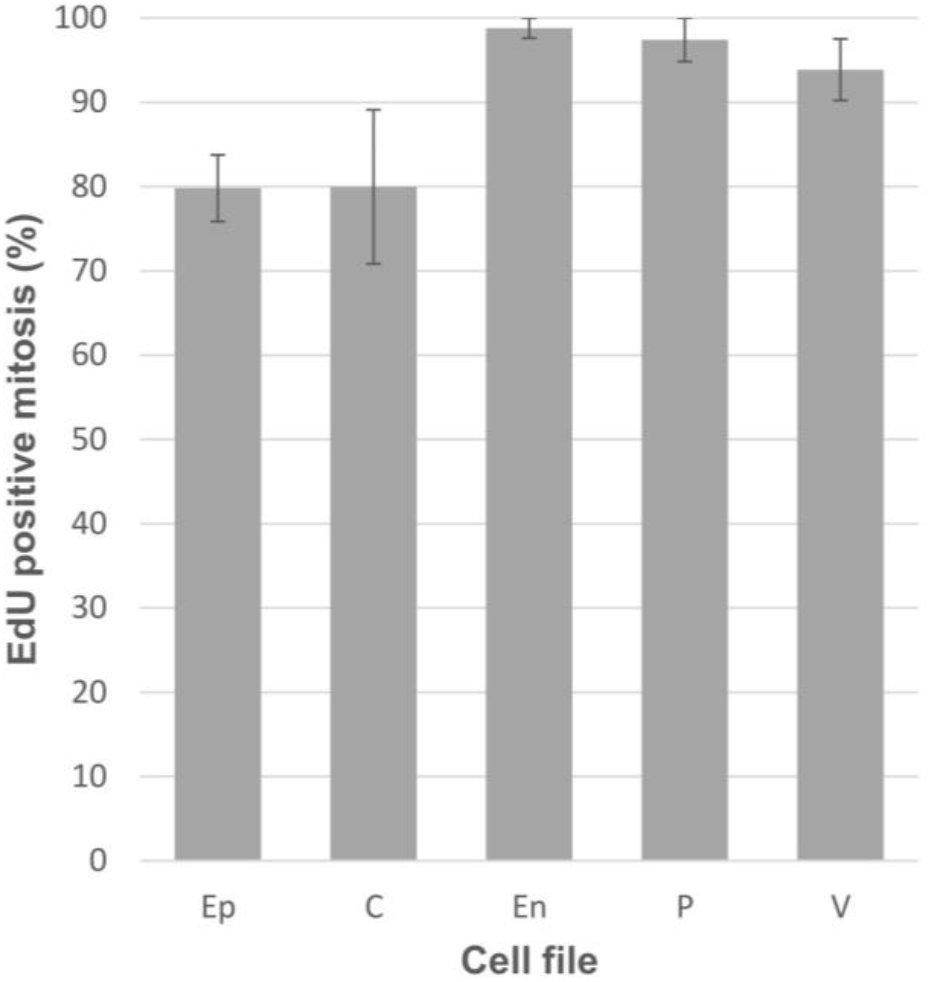
Average percent of EdU positive mitosis in different cell files. Five days old seedlings were incubated in the presence of EdU for 330 minutes, for the last 90 min colchicine was added. Seedlings were fixed immediately afterwards and subjected to analysis. Ep - epidermis, C - cortex, En - endodermis, P - pericycle, V - vasculature.

Considering gradients of the nuclei volume along the axis in the outer cell layers but not in the inner ones (Fig. 1), we speculate that the observed phenomena are related to differences in the chromatin condensation velocity at the end of G2 phase, which is in turn dependent on the nuclei volume.

G2 is the cell phase during which chromatin condensation to tightly packed chromosomes occurs (Pecinka et al., 2020). The velocity of this process seems to be directly dependent on chromatin shape and organization as visible in Figure 2. Dramatic differences in chromatin volume between pericycle and atrichoblast cells may led to differences in the kinetics of chromatin packing into chromosomes. In addition, increased volumes occupied by chromatin in the cortex and epidermis could be responsible for the extension of G2 duration in these cells, and seems to be dependent on the distance from QC.

### Mitosis duration

As an indirect measure of mitosis duration (including all stages), we employed the mitotic index, which reflects the relative duration of mitosis itself in relation to the whole cell cycle. We determined a mitotic index of 3-4% which means that the mitosis duration is noticeably short (about 20 - 25 minutes) and itself has only a limited contribution to total cell cycle duration.

### Determination of entire cell cycle duration

To investigate the entire cell cycle duration, we increased EdU incubation time to 8 hours. After this incubation period, gradients in EdU-positive cell frequencies from inner to the outer cell layers still exist. In the cortex cells 85% EdU positive cells have been detected in the proliferation zone, in contrast to the pericycle where it reaches 98%. In consequence, the entire cell cycle duration in the pericycle is less than 8 hours, while in the cortex and epidermis the process lasts up to 9 - 9.5 hours. These differences may be related to different chromatin landscapes, nuclei sizes, or even to nucleoli sizes, which can be detected in different cell files, even comparing trichoblast and atrichoblast cell (Figure S8). Unfortunately, it is exceedingly difficult to estimate the exact duration in outer cell layers because, in these layers, the duration correlates with the nuclear landscape (size) and chromatin status, which changes with QC’s distance (Figure 1). These data are in accordance with our observed differences in G2 duration and reflect differences in whole cell cycle duration as well. Interestingly, the EdU incorporation indices were significantly higher in the transition zone and reached 85% compared to 65% in the proliferation zone after about 5 hours of incubation (Figure S3). In consequence, we conclude that the duration of the endocycle is much shorter as the complete cell cycle in Arabidopsis.

**Figure S3.**
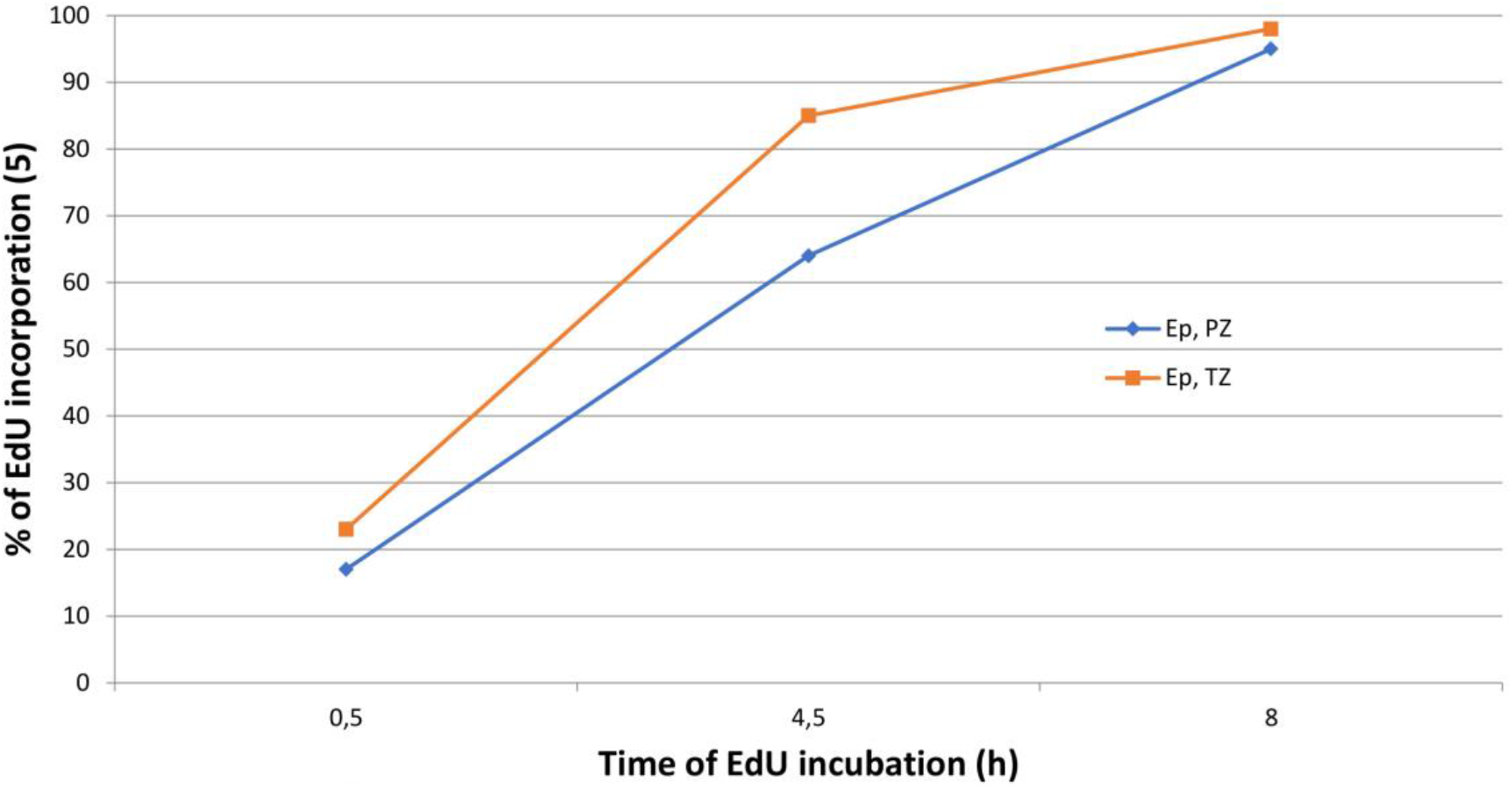
Ratio of EdU positive cell in the epidermis after 0.5; 4.5 and 8 h incubation in the presence of EdU. PZ-proliferation zone; TZ-transition zone (zone in which mitosis is not detected).

### Divergence in cell cycle duration after formative cell divisions

To further study potential differences in the root cell cycle duration, we analyzed cells with shared origin but different fate (Figure 6). Therefore, we investigated the ratio of cell production after tangential (in the epidermis) and periclinal (in the endodermis) cell divisions leading to the formation of daughter cells with different fates. The epidermis or rhizodermis of Arabidopsis thaliana roots is build up by usually eight cell files connected with two or more cortex cells (vertically) and about eight to sixteen cell files which connect with only one cortex cell. In 6 to 8-day old roots epidermal cells connected with two cortex cells divide tangentially and generate new cell file “continuities” which are connected with one cortex cell and develop later on to atrichoblasts (Berger et al., 1998). The daughter cells which are connected to one cortex cell produce approximately two times fewer daughter cells as cell connected to two cortex cells (Figure 6 D, E).

**Figure 6.**
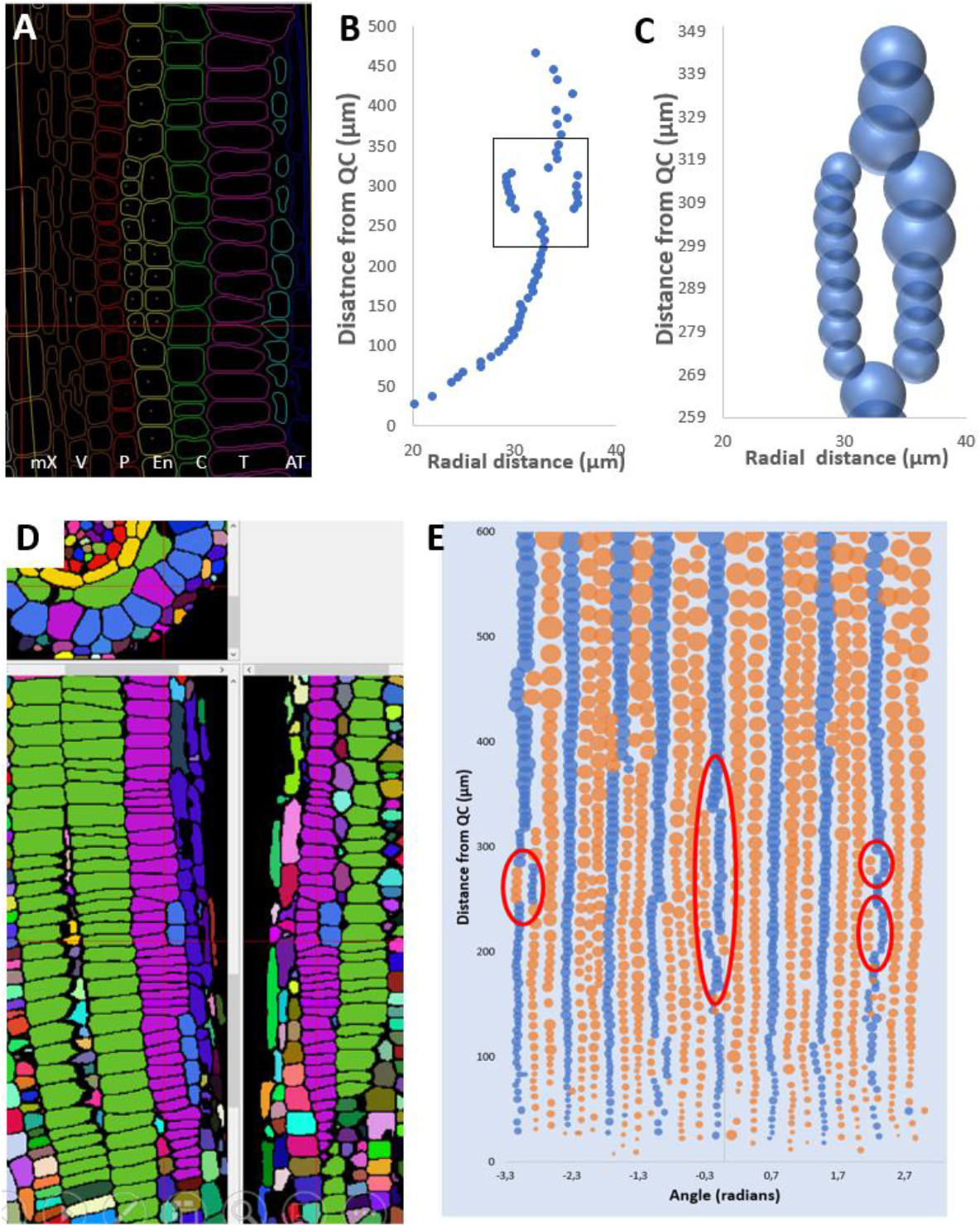
Differences in cell production rate after tangential and periclinal cell divisions. Roots of 7 days old seedlings were fixed, labelled for cell border detection, and segmented to analyse the precise tissue structures by iRoCS (Schmidt et al., 2014). Regions of interest related to periclinal (A, B, C) and tangential (D, E) cell division. Relative differences in cell volume are demonstrated by the size of bubbles (panel C, E). (A) mX (meta-xylem), V (vasculature), P (pericycle), En (endodermis), C (cortex), T (trichoblast precursor), AT (atrichoblast precursor); (E) - blue-Epidermal cells connected with two cortex cells; orange – epidermal cells connected with one cortex cell; red ovals show adjacent regions and cell files in the epidermis exhibiting differences in generated cell number.

In the endodermis periclinal divisions generate the formation of middle cortex and endodermis cells (Figure 6 A to C; Baum et al., 2002). Interestingly, the differences in the number of cells produced during the same time period are increasing with increasing distance from the QC (Fig. 6C). For example, after periclinal cell division in the endodermis, we have observed a lower cell production (fewer cell numbers) in the more outer cell files compared to the more inner ones. Interestingly, the fewer cells in the outer cell layer grow to bigger sizes in average – compensating for filling up the space within the organ. EdU-positive mitosis is more frequently observed (after shorter incubation time) in the cortex/epidermis cells with more proximal location to the QC, which means that cells with less proximal locations have a much longer G2 duration (Figure 4). In summary, these findings underline that cell cycle duration can be dependent on individual cell fate and position. An overview about average cell cycle duration and its phases in Arabidopsis is given in Table S1.

### Cell cycle duration in other species

In order to test the suitability of the presented approach in other species, we investigated, albeit in much less level of detail, roots of tomato, tobacco, and wheat (Figures S4-S6).

**Figure S4.**
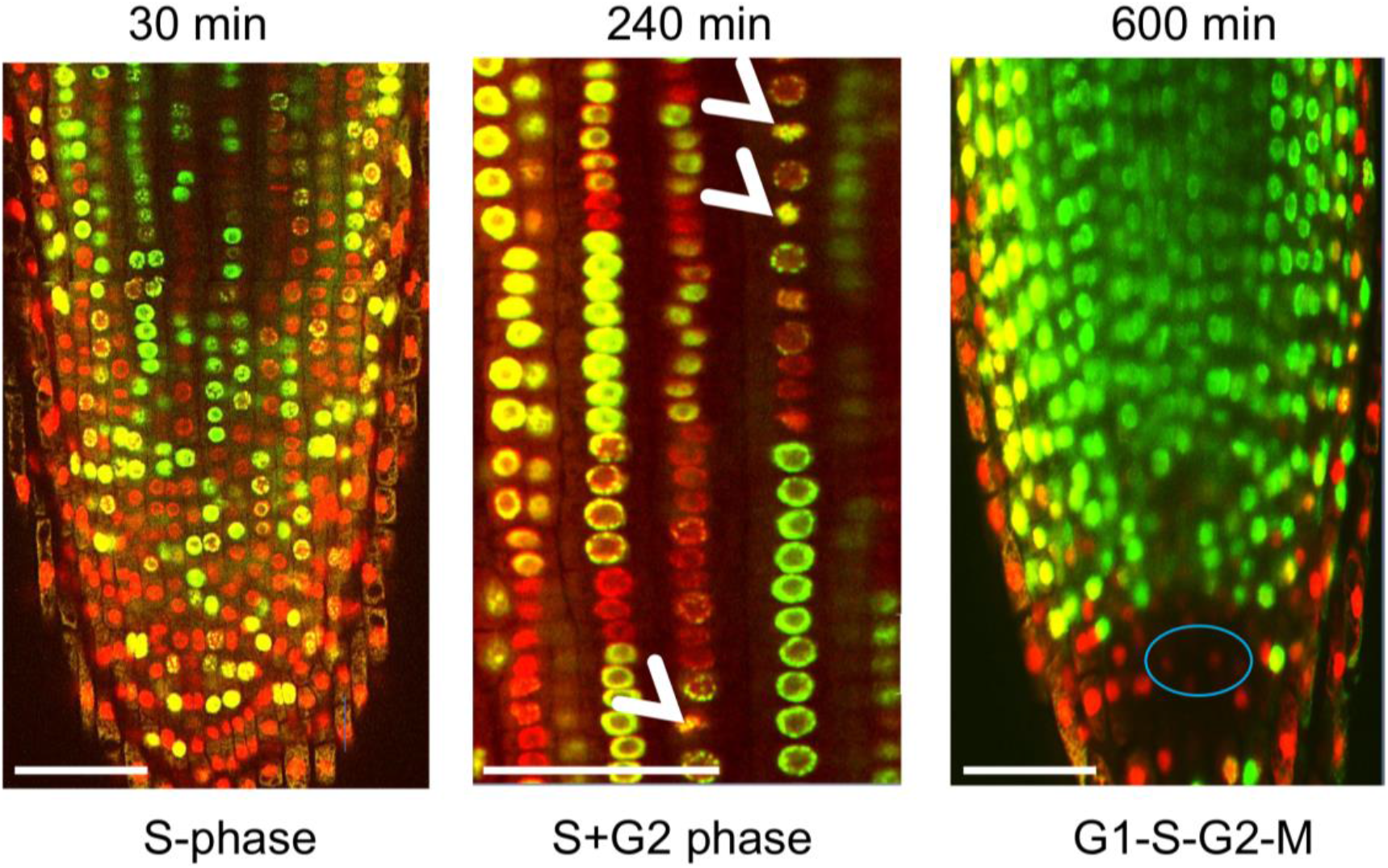
Cell cycle analysis in tomato root. 4-5 days old seedlings were incubated with EdU for 30 min, 240 min, and 600 min. 30 minutes of incubation showed entire colocalization DAPI and EdU on cortex (2-3) focal plane; 240 min incubation led to the first appearance of EdU-positive mitosis (white arrows) and 600 min incubation to distinguish the stem cell niche. Nuclei are depicted in red; EdU positive nuclei are shown in green; the QC region (blue oval) is very dense and shows weak DAPI staining. Scale bar-100 µm.

**Figure S5:**
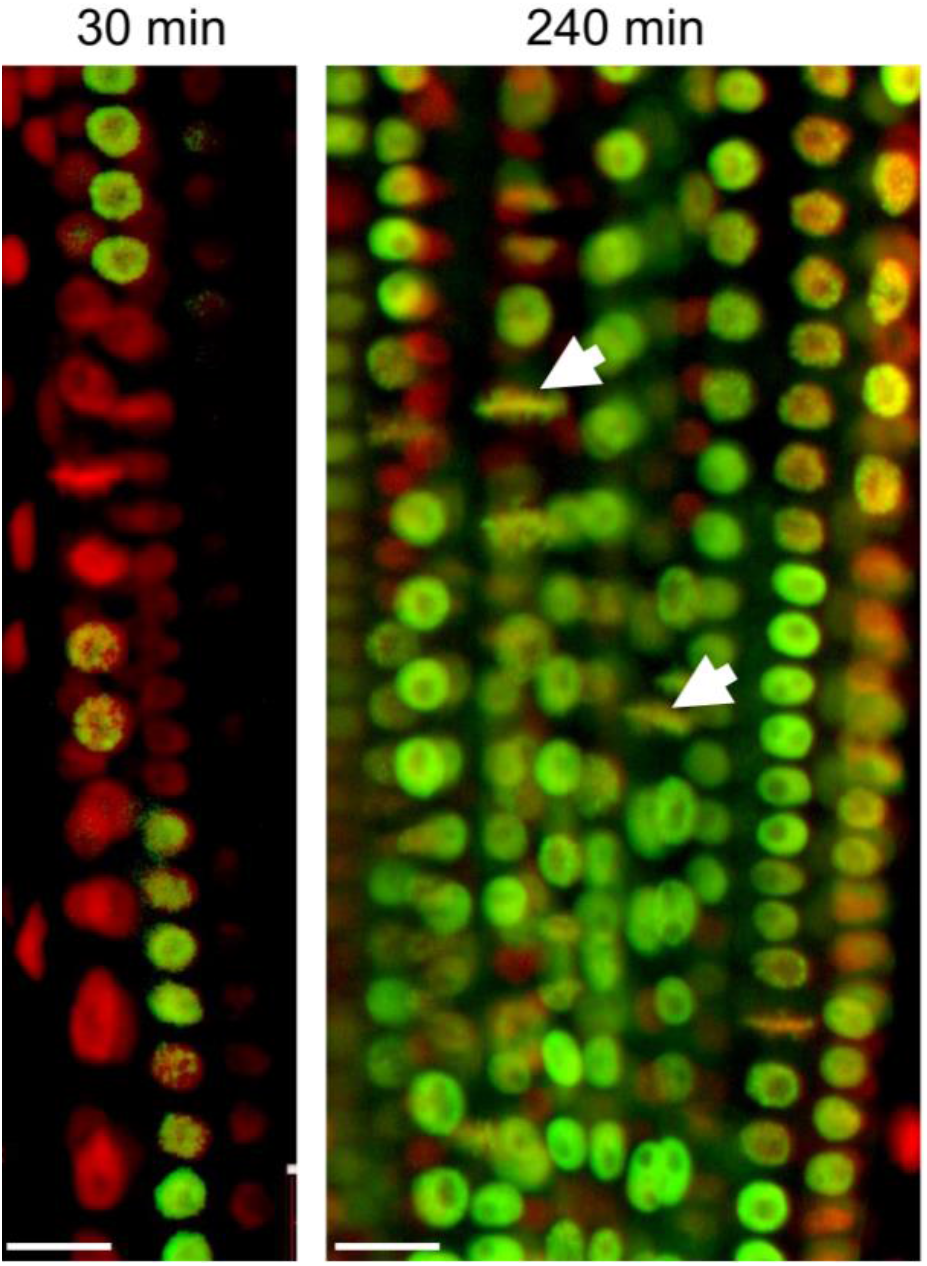
Cell cycle analysis in tobacco. 4-5 days old seedlings were incubated with EdU for 30 min and 240 min. Arrow show selected mitotic plates. Nuclei are depicted in red; EdU positive nuclei are shown in green; Scale bar-20 µm.

**Figure S6:**
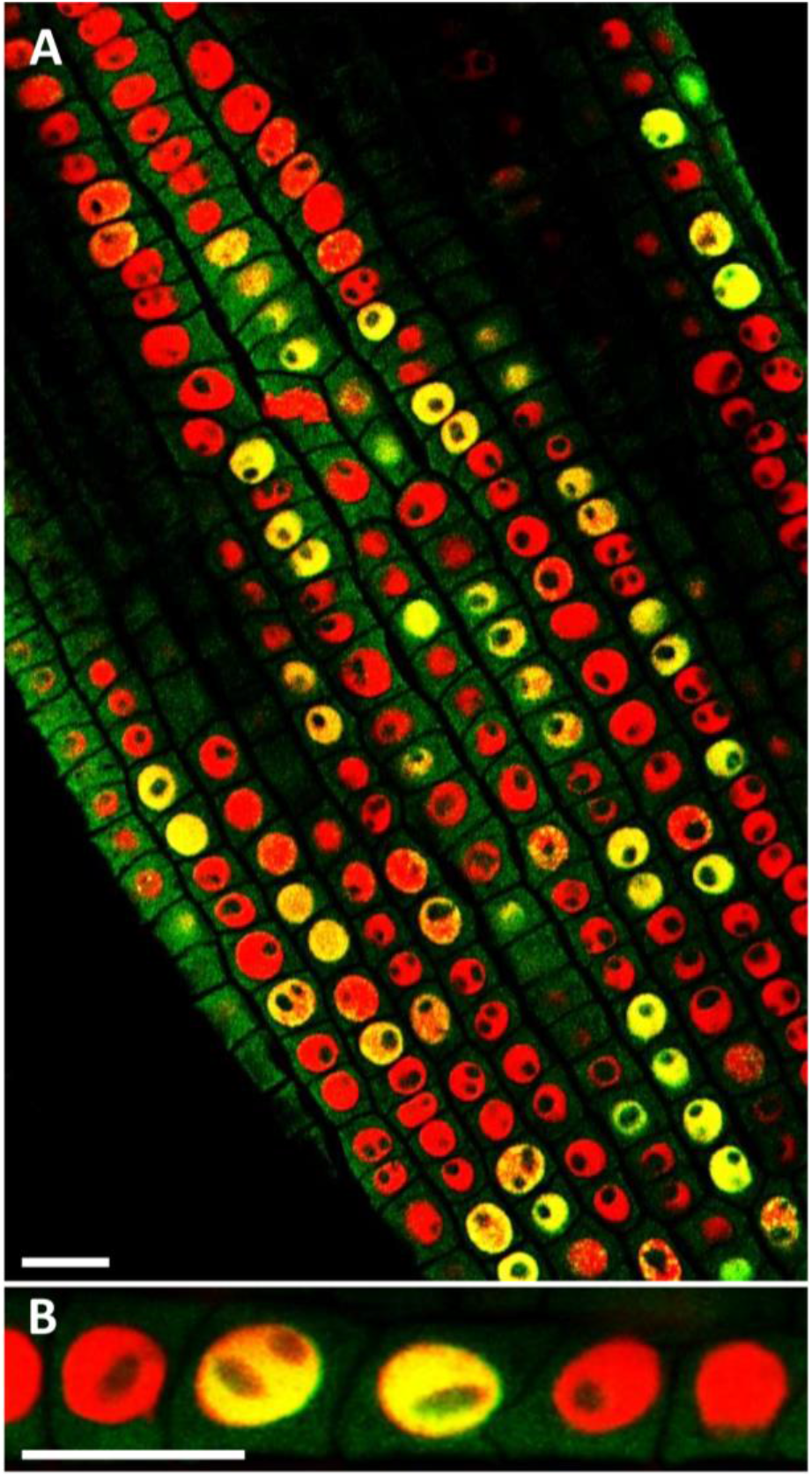
EdU incorporation into wheat tissue. Four days old wheat seedlings were incubated with EdU for 35min. (A) – general view; (B) – close up view. Overlay images, DAPI stain of nuclei is red coloured, EdU signals are coded in green. Scale bar-20 µm.

For S-phase duration we detected EdU incorporation with almost full DAPI co-localization after 20-35 minutes of incubation in these three species. EdU-positive mitotic events plates were detected in tomato and tobacco after 240 minutes of incubation. Interestingly, in tobacco and tomato, no obvious differences in nucleus/chromatin organization between outer and inner cell layers were detected. In consequence, in these species, very similar G2 durations occur in all cell files analysed. Additionally, in tobacco and tomato, the duration of DNA replication and G2 phases are even shorter than Arabidopsis. Moreover, in the case of tomato, almost all cells became “EdU-positive” after 9-10 hours of incubation except in the stem cell niche where cells may be less accessible for analysis due to enhanced density and attenuated stain accessibility (Figure S4).

## Discussion

Plant organ growth is regulated by a complex and dynamic set of multifactorial events that need multi-scale approaches to identify different parameters for precise coordination between different cell types, individual cell position, rate of cell production, and cell expansion.

Even a simple organ like an Arabidopsis root possesses significant divergence of processes controlling geometry of cells and organs and the cellular chromatin status, which eventually controls the subtle differences in cell cycle kinetics in the different files. In addition to cell volume, the chromatin organization may have a direct impact on the cell cycle. From the geometrical point of view, cell cycle transitions can be considered as changes in chromatin status (Ma et al., 2015). Nucleosomes and the associated chromatin structure must disassemble before replication but re-assemble afterwards (Margueron and Reinberg, 2010) while significant chromatin condensation occurs during mitosis. Therefore, it seems logical that both transitions were dependent on chromatin geometrical organization: compact nuclei like QC nuclei exhibit a delay in DNA replication, while nuclei with diffuse chromatin (cortex/epidermis) may have a delay in chromosome condensation (extended G2 duration).

### Cell culture-based methods

Classically cell culture based methods have been dominating the field of cell cycle analysis as they offer advantages of sampling nearly homogenous population of cells (Gould, 1984). But unfortunately, this approach is limited because it neglects potential positional effects on cell cycle kinetics regarding interactions of individual cells within intact tissues and organs what led to differences in chromatin landscape. But still important features have been learned by cell culture techniques such as the relative durations of each cell cycle phase, elucidation G1:S:G2 ratios of 50:10:40, revealing that the S-phase represent 5-10% of cell cycle duration, and that the G1 phase is roughly 20% longer than G2 (Yanpaisan, W., et al., 1998; Menges, M., & Murray, J. A., 2002). Additionally, these methods allowed to identify cell cycle genes and their functions which led to a simplified concept of main hubs regulating root development but missing potential roles of cell file gradients or changes in chromatin organization as critical regulatory mechanisms (Desvoyes et al., 2014; Otero et al., 2016).

### Duration of DNA replication

Many methods allow monitoring DNA replication and hence S-phase duration by using markers such as H^3^-thymidine and more recently BrDU and EdU (Mickelson-Young, L.et al., 2016; Pereira P. et al., 2017). These studies claim that DNA replication duration spans about 2.5-3 hours based on estimation of the time after which incorporated marker fluorescence intensities reaching a maximum. However, substitution of regular growth media with EdU containing media without pre-adaptation before onset of analysis may distort precise cell cycle progression results. When cellular thymidine pools are high, EdU will incorporate in parallel with thymidine potentially leading to an overestimation of cell cycle duration (Hayashi et al., 2013). Therefore, new protocols should consider potential consequences of immediate medium exchange as proposed in the majority of published protocols (Van’t Hof J. 1967; 1969). Consequently, in our protocol, we propose root adaptation to the new medium to prevent potential disturbances in determining cell cycle durations.

An indirect method to estimate the duration of DNA replication is to determine the number of cells in S-phase by flow cytometry in not-synchronized suspension cells. It was found that the ratio of cells which undergo DNA replication is not exceeding 5-7% of all cells analysed. In consequence, the period of DNA replication does not exceed 5-7% of the total cell cycle duration.

### Molecular marker lines for quantitative analysis

Another popular route is the use of molecular markers to investigate cell cycle dynamics in living roots (Yin K. et al., 2014). Despite the fact that these markers more or less precisely reflect cell cycle kinetics major problems may occur when markers like CyCB1.1::GFP are less well detectable in inner cell layers like pericycle. Additionally, their expression reflects rather the G2 phase, but not the mitosis itself (Lavrekha et al., 2017). Another limitation of its general applicability is that marker lines for cell cycle events have to be generated *de novo* for each plant species of interest as well as for Arabidopsis mutant analysis. To overcome these time-consuming transgenic procedures Bhosale et al. (2018) have recently proposed to employ DNA staining techniques for analysis of DNA replication. Another protocol based on multicolor *in vivo* markers for each cell cycle stages was recently applied to monitor Arabidopsis root cells cycle progression features (Desvoyes et al., 2020). But all *in vivo* methods using fluorescing markers should take into account potential effects of long-term laser irradiations during microscopic data acquisition on the cell cycle dynamics itself (Tosheva et al., 2020).

### Positional cell cycle gradient analysis

Our method combines and extends all of these features by simultaneously performing a detailed analysis of all cell cycle events in each cell file independently and simultaneously using chemical labeling, high-resolution confocal imaging, and subsequently rigorous, quantitative image analysis. For this we first optimized application and timing of EdU incorporation as a marker for DNA replication by exposing Arabidopsis seedlings to EdU for different time periods (Figure 3) and determined the distribution of EdU-incorporation in all root cells simultaneously. Then we applied the iRoCS pipeline for 3D-high-resolution analysis of the root organ (Schmidt et al., 2014). As shown previously this analysis is not limited to Arabidopsis, for which it has been developed at first, but can easily be adopted to other plants as well (Pasternak et al., 2017). Using this pipeline, which resolves all root cells and root cell files simultaneously, we found already after 20 min exposure EdU positive cells in the proliferation domain as well as in the transition zone, the domain where cells stop to divide and start to slightly elongate (Pasternak et al., 2017; Lavrekha et al., 2017) but undergo endo-reduplication. Longer incubation times of up to 4 hours allowed to detect EdU-positive mitotic cells undergoing DNA replication, moving into G2 and entering into mitosis. By consequent changing the EdU-incubation time we could use the distribution of EdU-positive mitotic events to quantify S+G2+M duration. For example, 5 hours after EdU incubation, we found that almost all elongating cells performed replication, which means that all cells in this zone passed G1-S transition. This allowed us to conclude that the entire cycle duration in this region is around five hours (Figure S3,, S7).

**Figure S7.**
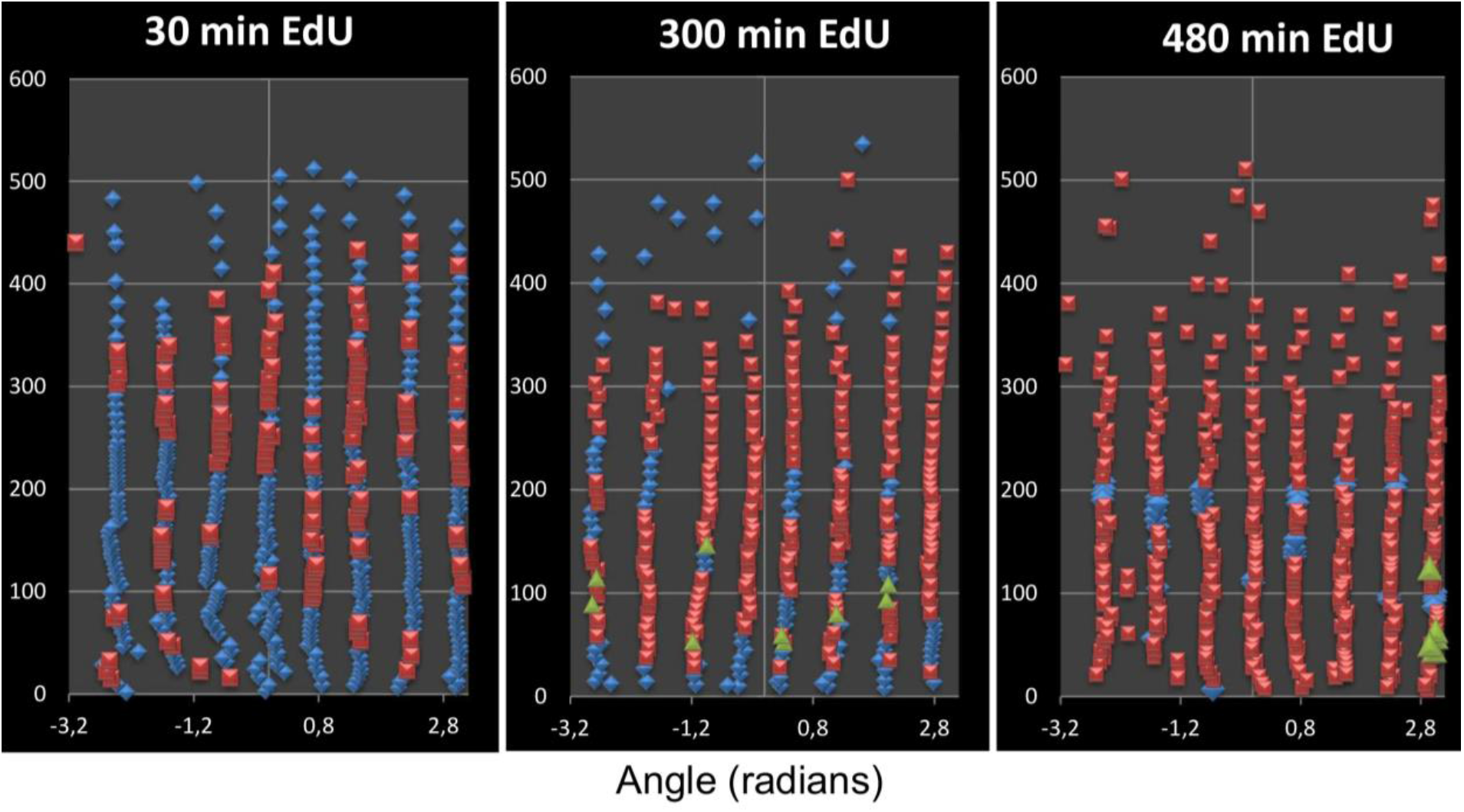
EdU incorporation into cortex cells after incubation for 0,5; 5 and 8h. Cells without EdU are depicted in blue, EdU-positive cells in red, and mitotic cells in green.

Overall, our method allows detailed analysis of all cell cycle events in each cell file independently and simultaneously. Previously, we demonstrated significant differences in cell geometry and sizes (Blein et al., 2018) among different cell layers what should eventually involve differences in cell cycle duration for harmonized organ growth. Here, we were able to describe significant differences in cell cycle duration of cells with different fate and show that cell cycle duration is much slower in the cortex and epidermis compared to pericycle and is accompanied by a progressive slowdown of their cell cycle as a function of QC’s distance. Such differences may serve to compensate faster increases in cell volume in outer cell layers (Figure S1; see Blein et al., 2018). Moreover, this difference is directly related to nucleus geometry: a significant increase in nuclei volume in parallel to cell volume led to an extension of the time required for chromosome condensation. Finally, we observed differential cell cycle durations after tangential or periclinal cell divisions leading to cell lines with different fates Thus, our study provides direct evidence for dynamic positionally driven regulatory mechanisms controlling cell cycle duration.

A comparative summary of the different methods is given in Table S2.

**Figure S8.**
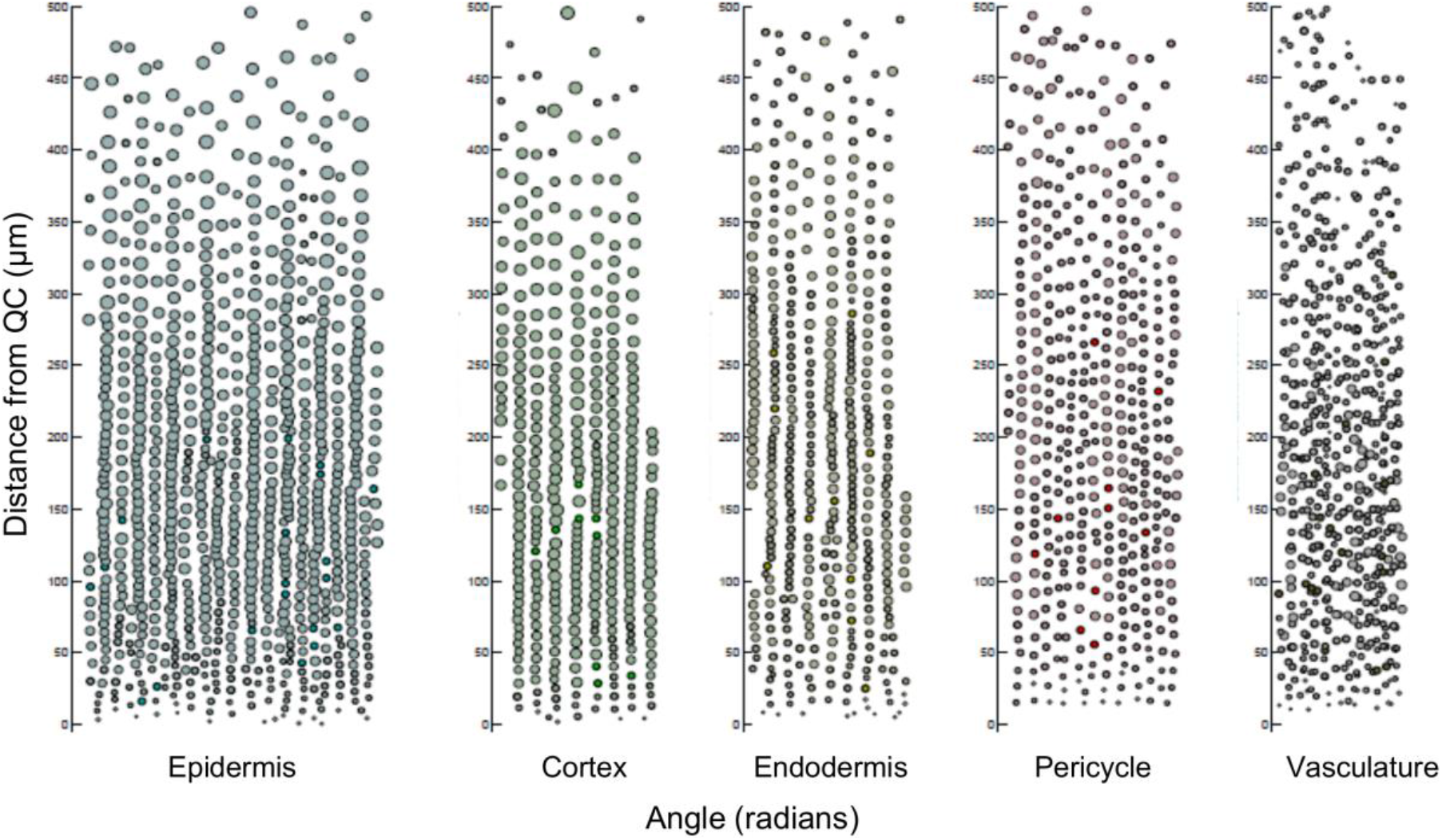
Distribution of the nucleoli volume along the root axis in different cell files. Volumes of nucleoli in each cells of the root were determined. Y-axis-distance from QC; X-axis-radians; Nucleoli volumes are proportional to area of the circle.

## Conclusions

Root growth is a dynamic process and occurs in a strictly coordinated manner. Consequently, dynamic changes in cell elongation, volume, cell proliferation of involved tissues must be strictly balanced. Here we present a marker-free and straightforward method for rapid and simultaneous estimation of the cell cycle duration and its stages in each cell file of plant roots. Furthermore, the method does not require significant mathematical calculations. Using this high-resolution pipeline, we show that differences in the duration of cell cycle stages correlate with differential cell volume kinetics, the number of cells in distinct cell files, and their nuclear landscape. Additionally, the method is not restricted to Arabidopsis but is also applicable to other plant species. From our data, we conclude that roots keep their structural integrity at least partially by differential cell production rates reflected by distinct cell cycle duration timing.

## Materials and methods

### Plant materials and growth conditions

*Arabidopsis thaliana* (L.) Heyhn. ecotype Col-0, *Nicotiana tabacum cv*. Samsun N.N., *Lycopersicum esculentum* L., and *Triticum aestivum* L. were used in experiments. Arabidopsis seeds sown on square Petri dishes containing TK1 medium (Pasternak et al., 2020). The plates were kept at room temperature for 4 h before being transferred to 4°C for 12 h. The plates were then transferred to 22°C under a 16 h / 8 h light/dark photoperiod with a light intensity of 80 µmol·m^-2^·s^-1^ for 4.5 days. The seedlings were transferred to a 6-well plate containing liquid TK1 medium. After a 12 h incubation in similar conditions as on agar medium growth conditions, 2.5 mM colchicine and 10 µM EdU (5-ethynyl-2’-deoxyuridine) were added at defined time points (Figure 3) and seedlings from all vials were fixed 8 h after onset of first EdU addition. Seeds of the *Nicotiana tabacum* Samsun NN and *Lycopersicum esculentum* L. were surface-sterilized in 0.2% NaClO and sown on square Petri dishes containing T1 medium (supplemented with 2% sucrose) optimized for root growth (Pasternak *et al*., *1999)*. 4-5 days old seedlings have been used for labelling. Wheat seeds were soaked in water in Petri plates. 3-5 days old seedlings were transferred to liquid T1 medium for 16 h adaptation.

### Pipeline for determination of cell cycle duration

For determination of cell cycle duration 10 µM EdU were added to the seedlings in the liquid medium 12, 14, 16, 18.5 and 19.65 hours after starting of adaptation for each plate separately. Seedlings were fixed 20 h after onset of incubation. This timeline allowed us to apply different EdU-incubation times for either 8 h; 6 h; 4.5 h; 1.5 h or 20 min. For determination of (G2+S) duration colchicine was added to the plate after 180 and 270 min EdU incubation. These plants were fixed after further incubation for 90 minutes (see Figure 3 for experiment design).

### Staining

After EdU incubation plants were fixed in microtubule stabilizing buffer containing 4% formaldehyde (MTSB) and EdU incorporation was detected as described previously (Pasternak et al., 2015).

For volume determination, roots were fixed and labeled with the modified Truernit et al. (2008) method. Scanning and analysis procedures were done essentially as described in Schmidt et al., 2014; Pasternak et al., 2021.

### Confocal imaging

DAPI/EdU-stained samples were recorded using a confocal laser scanning microscope (LSM 510 META NLO; Carl Zeiss, Oberkochen, Germany) with a LD LCI-Plan-Apochromat 25x/0.8 DIC Imm Corr objective. For DAPI excitation, a Chameleon laser adjusted to 740 nm excitation was used and emission was detected with a band pass filter (BP 390-465 IR). EdU excitation was at 488 nm and emission detected with a band pass filter (BP 500–550 IR). PI-stained roots were recorded using a confocal laser scanning microscope (ZEISS LSM 510 META NLO) with a LD LCI-Plan-Apochromat 25x/0.8 DIC Imm Korr objective with glycerol as immersion medium. Excitation wavelength was 543 nm and emission was detected with a long pass filter (LP560). Serial optical sections were reconstituted into 3D image stacks to a depth of 250 lm with in-plane voxel extents of 0.15 lm and a section spacing of 0.9 lm in the z-direction. Alternatively, all other suitable microscopes equipped with a diode 405 nm laser can be used. Serial optical sections were reconstituted into 3D image stacks with in-plane (x-y) voxel extents of 0.15 μm and 0.9 μm section spacing (z). Three to five overlapping images (tiles) were recorded for each root.

### Image processing and analysis

Image analysis were performed essentially according to Pasternak et al., 2021. Briefly, images were converted to hdf5 format using the LOCI plugin for ImageJ (http://imagej.nih.gov/ij), then stitched together to obtain a root tip total length of 400 μm from the QC using xuvTools (Emmenlauer et al., 2009). Finally, 5-10 representative roots were chosen for detailed annotation. The DAPI and EdU channel images were processed with the iRoCS Toolbox (Schmidt et al., 2014) in the following way: nuclei were automatically detected using the “01-Detect Nuclei” plugin, then the epidermis was semi-automatically labelled using the “02-Label Epidermis” plugin. After the QC position was marked (Channel->New Annotation Channel), the nuclei were set in spherical coordinates using the “03-Attach iRoCS” plugin. Automatic classification of the nuclei to the corresponding cell types (epidermis, endodermis, cortex, pericycle, vasculature, root cap) was done using the “04-Assign Layers” plugin, which also enabled the automatic annotation of nuclei in mitotic state (option “Re-classify mitotic state”). All annotated roots were manually corrected for erroneous layer, mitosis, and EdU assignments.

For the nucleus geometry analysis, original nuclei-labeled images were converted to .tiff, and OTZU methods were applied. Thereafter nucleus edges have been detected, images were converted to hdf5 and further segmentation was performed accordingly to the standard pipeline (Pasternak et al., 2021).

### Data analysis

The data processed by iRoCS were exported to .csv files and analyzed with Microsoft Excel. Statistical analysis was done using Student’s *t*-test. Roots were virtually divided to 50 µm sections. The DNA replication index (DRI) was calculated as the proportion of cells posing DNA replication during incubation time to all cells in the current section and was calculated for each 50 µm interval.

## Acknowledgements

We thank Dr. Roland Nitschke and the staff of the Life Imaging Center (LIC) in the Center for Biological Systems Analysis (ZBSA) of the Albert-Ludwigs-Universität Freiburg for help with their confocal microscopy resources, and the excellent support in image recording and analysis. Experimental work was supported by Bundesministerium für Bildung und Forschung (BMBF SYSBRA, SYSTEC, Haploswitch), the Excellence Initiative of the German Federal and State Governments (EXC 294), DFG (INST 39/839-1 FUGG), SFB746 and Deutsches Zentrum für Luft und Raumfahrt (DLR 50WB1022) as well as a DFG grant (KI 1077/2-1) to SK.

## Author Contributions

Conceptualization: TP, SK, KP; methodology: TP, Sample preparation, scanning, image processing: TP; resources: KP; data curation: TP, SK, KP writing: TP, SK, KP. All authors have read and agreed to the published version of the manuscript.

---

**Table S1.** Summary of cell cycle duration in Arabidopsis RAM.

| Duration of the EdU incubation | Cell cycle stage |  |
| --- | --- | --- |
| 20-30 min | DNA replication | Cell cycle/endocycle |
| 3-5 hours | S-G2-M |  |
| 8-10 hours | S-G2-M-G1 | Entire cycle duration |
| 18-24 h | S-G2-M-G1 | Status of the stem cell niche |

**Table S2.** Comparison of different methods for cell cycle duration estimation.

| Method<br>Authors<br>Parameters<br>extracted | H<sup>3</sup>thymidine<br>/colchicine<br>Clowes<br>1961 | H<sup>3</sup>thymidine<br>/colchicine<br>Van't Hof<br>1967 | EdU Hayashi et<br>al.,2013 | Kinematic<br>Ivanov/Dybrowsky 1997 | *In vivo*<br>Method<br>Yin et al.,<br>2014 | Stripflow<br>Yang et al.,<br>2017 | „Plant cell<br>cycle<br>indicator“<br>Desvoyes et<br>al., 2020 | This<br>paper |
| --- | --- | --- | --- | --- | --- | --- | --- | --- |
| Cell<br>type/subtype/position | Yes | no | partially | only cortex | partially | only cortex | no | yes |
| Direct detection | Yes | yes | yes | no | partially | no | yes | yes |
| Link to chromatin<br>structure | No | partially | no | no | yes | no | no | yes |
| Pre-adaptation | No | no | no | yes | yes | yes | yes | yes |
| Marker introduction<br>to the mutant line | no | no | no | no | 3-5 months | no | 3-5 months | no |
| Cell cycle stages<br>resolved | Partially | partially | no | no | yes | no | yes | yes |

